## Supplementary Information for "Inadequate structural constraint on Fab approach rather than paratope elicitation limits HIV-1 MPER vaccine utility"

List of content:

Supplementary Figures 1 to 15

Supplementary Tables 1 to 5

References

**Supplemental figures**

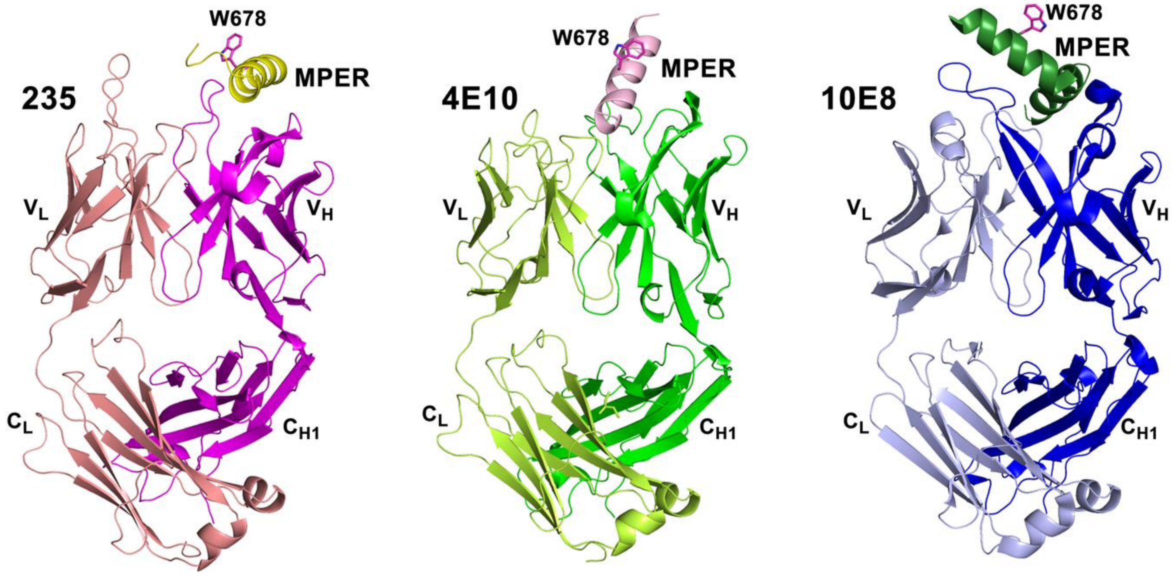


**Supplementary Fig. 1**. **Ribbon diagrams of three Fabs targeting the MPER C-terminal helix in complex with MPER peptides.** Fabs 235 (PDB: 8FYM), 4E10 (PDB: 2FX7) and 10E8 (PDB: 4G6F) are displayed as ribbon diagrams and placed in a similar orientation. MPER residue W678 is shown in stick representation to indicate the relative orientation of the MPER C-terminal helix in each structure.

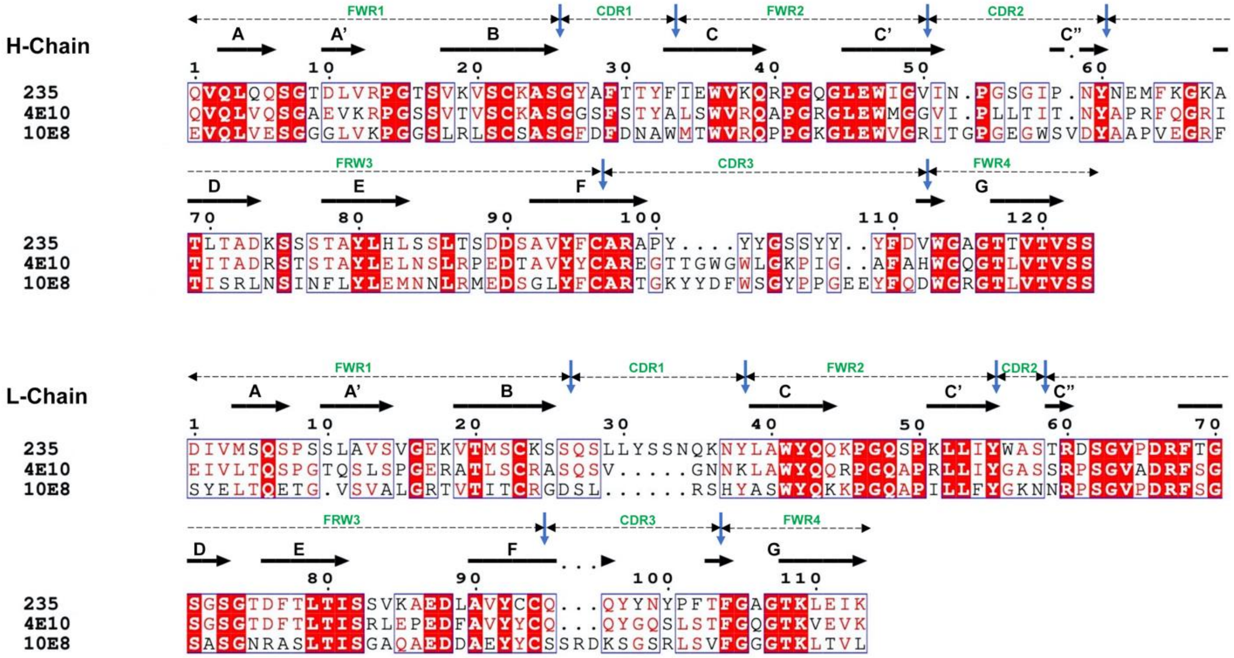

**Supplementary Fig. 2**. **Structure-based sequence alignment of variable domains of Fabs 235, 4E10 and 10E8**. β-strands are marked with arrows above the appropriate sequences of Fab235. There is a bulge (not shown) in the G-strand of both the H- and L-chain. The complementarity-determining regions (CDRs) and framework regions (FWRs) are labeled in green. Identical residues are highlighted in red. The alignment between mouse 235 and human bnAbs was generated with the programs MultAlin (1) and ESPript (2) and modified for clarity. Note that germline genes of VH domain segments (V, D and J) for Fab235 are IGHV1-46*01, IGHD4-23*01 and IGHJ2*01, and IGKV4-1*01 and IGKJ4*02 for the VL domain segments.

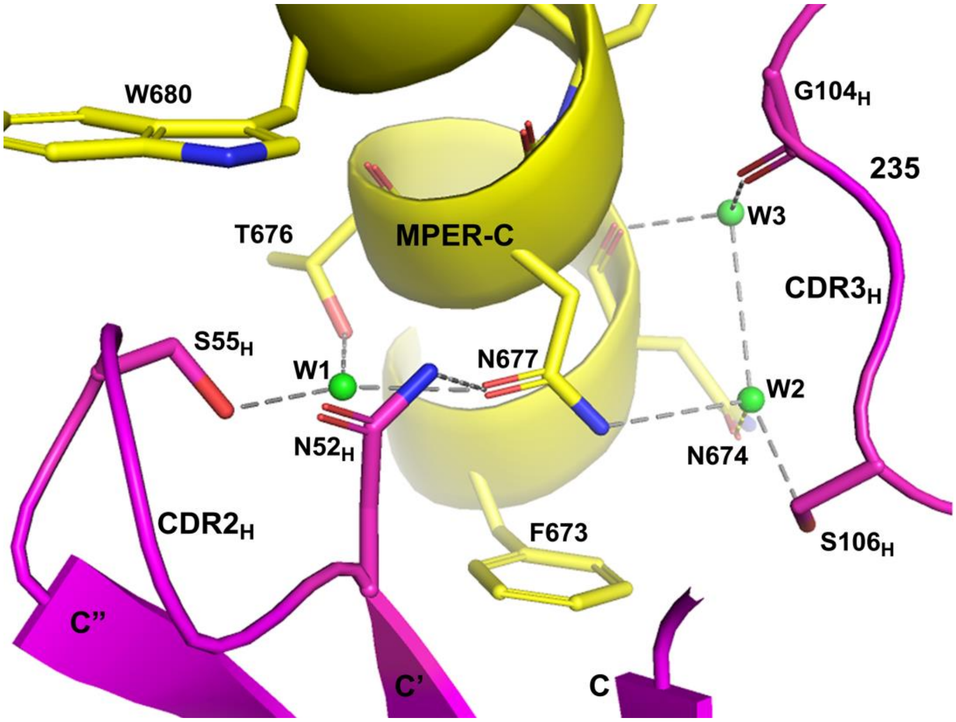


**Supplementary Fig. 3. The water-mediated hydrogen-bond network featured in the interaction of Fab235 with MPER-C.** Water molecules are depicted as green spheres and labeled W1-W3. Water molecule W1 is conserved in all four complexes in the asymmetric unit of the crystal. In Fab235/MPER-C complexes DEF and GIJ, water W2 appears to be replaced by a chlorine ion and water W3 is absent.

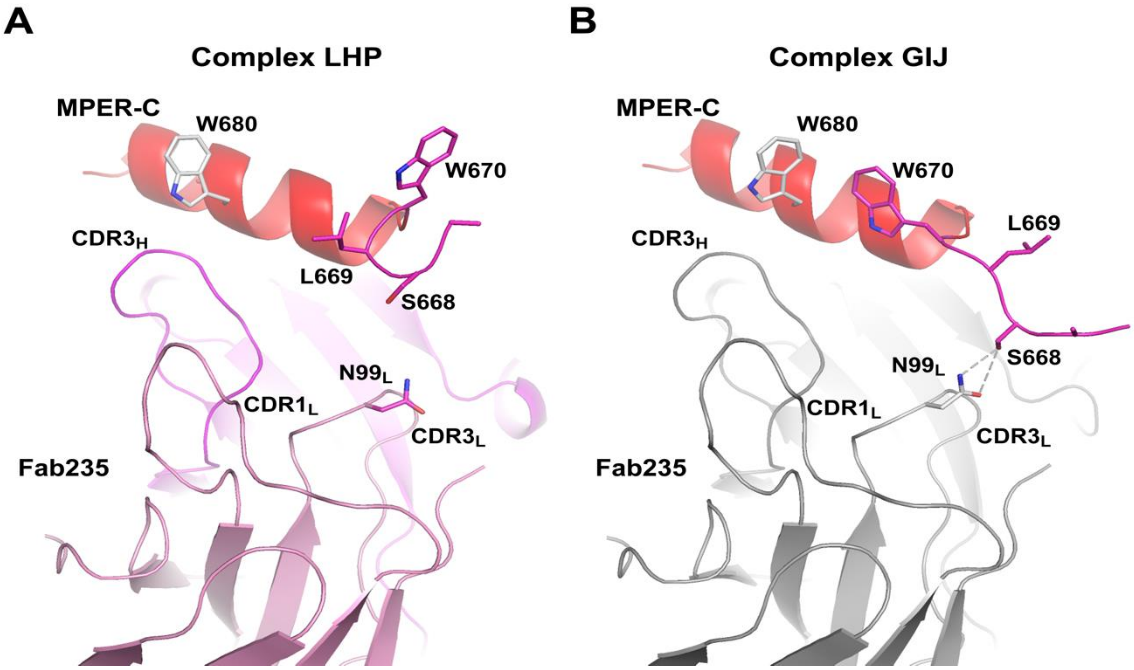


**Supplementary Fig. 4. Comparison of the four Fab235-MPER-C complexes in the asymmetric unit of the crystal reveals the conformational dynamics of the N-terminal loop region of MPER-C and its potential interaction with Fab235.** (**A**) Representative conformation of the loop region found in complexes LHP and ABC. (**B**) Representative conformation of the loop region found in complexes DEF and GIJ and its interaction with Fab235. The latter two complexes are characterized by a hydrogen bond between S668 and $N99_L$. Given the impact of the S668A mutation on Fab binding (Fig. 1D), it is likely that the two conformations can interchange.

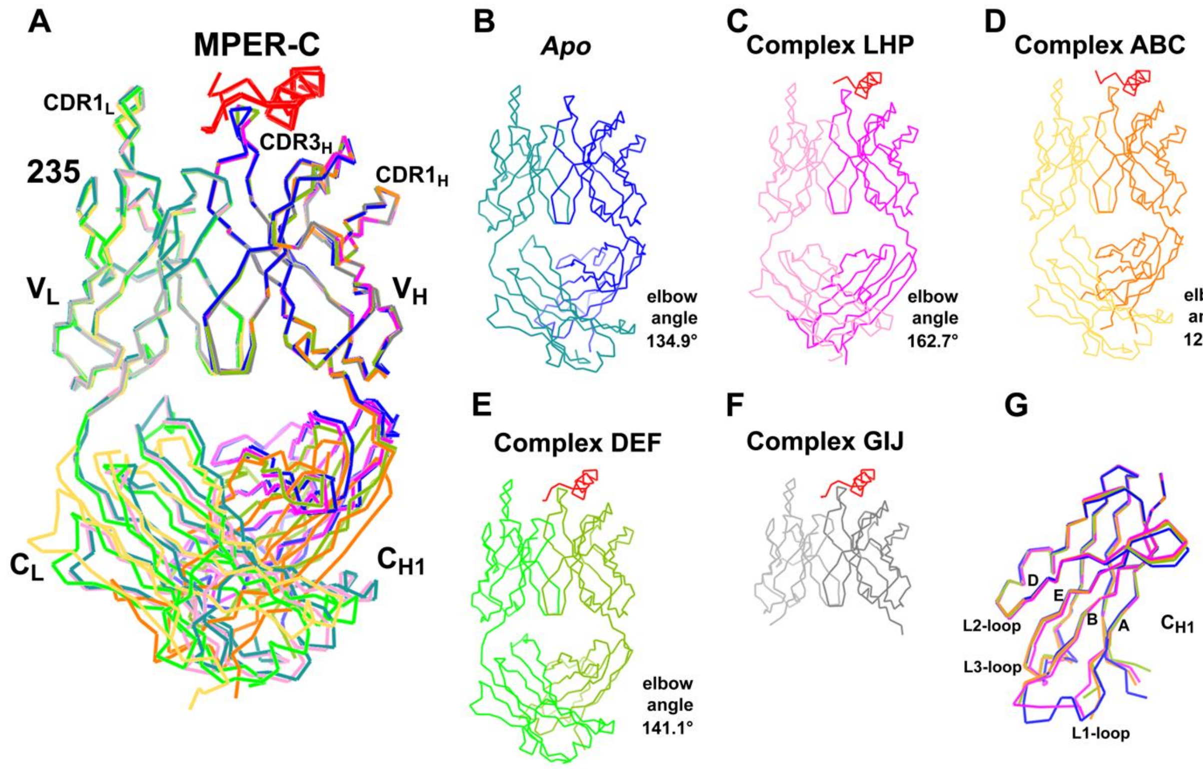


**Supplementary Figure 5. Comparison of the structures of Fab235 alone and in complex with MPER-C.** (**A**) Least-square alignment of the structures of Fab235 alone and the four structures of Fab235 in complex with MPER-C based on their $V_H$ domains. The structural alignment indicates minimal, if any, conformational change upon MPER-C binding either within individual $V_H$ or $V_L$ domain or between these two V domains. As shown, subtle changes were observed in $CDR1_H$, $CDR3_H$ and $CDR1_L$, even though $CDR1_L$ was not even involved in the interaction with MPER-C. Colors are the same as those used for the individual structures shown in panels B-F. (**B-F**) C$\alpha$ trace representations of Fab235 alone and in the four complexes with MPER-C, with elbow angles denoted. The C domains of complex GIJ are completely disordered and hence not visualized. (**G**) The structures of the Fab235 $C_{H1}$ domains from the *Apo* structure and the three complex structures (omitting complex GIJ) were aligned for comparison of their three bottom loops. The A-, B-, D- and E-strands of the domain are labeled for easy identification of the locations of these loops. The colors of the H- and L-chains in panels B-G are the same as in panel A.

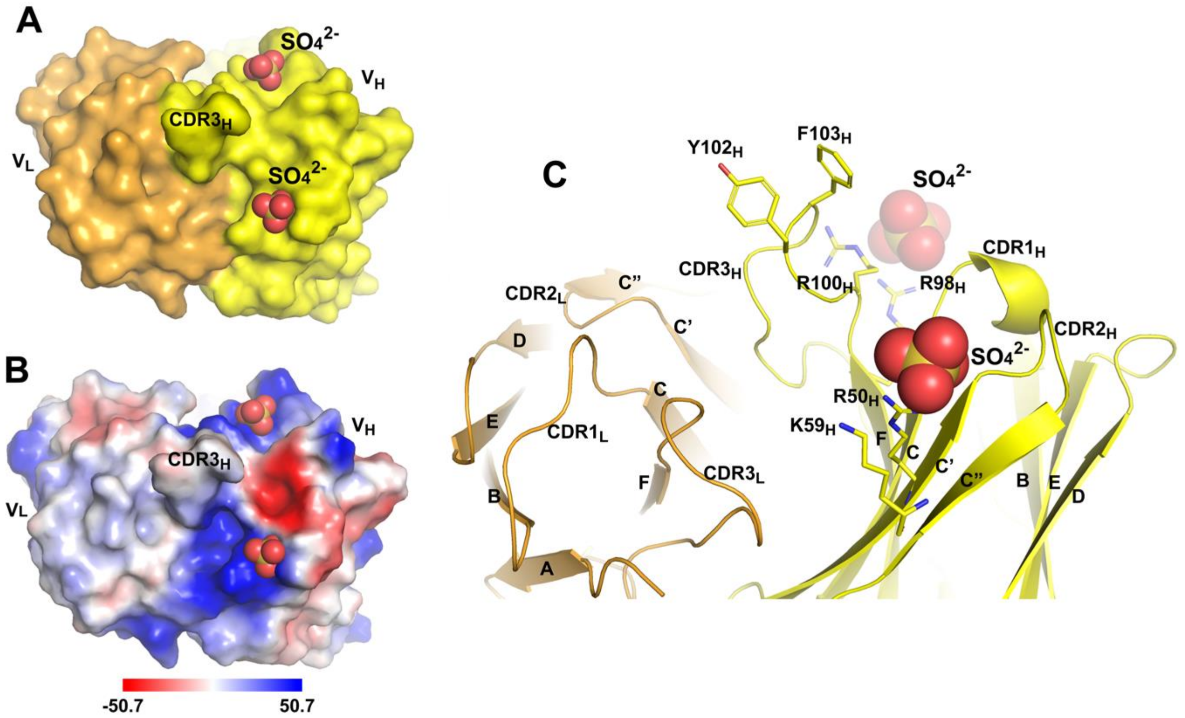


**Supplementary Figure 6**. **CDRs of Fab460.** (**A**) Top view of a surface representation of the Fab460 CDRs. The heavy and light chains are colored in yellow and orange, respectively. Two sulphate anion groups bound in the structure are shown in sphere representation. The tip of $CDR3_H$ is labeled. (**B**) Top view of Fab460 CDRs shown in electrostatic potential surface representation ranging from positive (red) to negative (blue) with scale range utilized below. The heavy chain CDRs form charged patches or pockets. Two sulphate anion groups from the crystallization buffer are trapped in positively charged pockets. (**C**) Side view of the Fab460 CDRs shown in ribbon representation. Residues $Y102_H$ and $F103_H$ protrude from the tip of $CDR3_H$ and are shown as sticks. Residues $R50_H$ and $K59_H$ and $R98_H$ and $R100_H$ contribute to the formation of two positively charged pockets and are also shown as sticks.

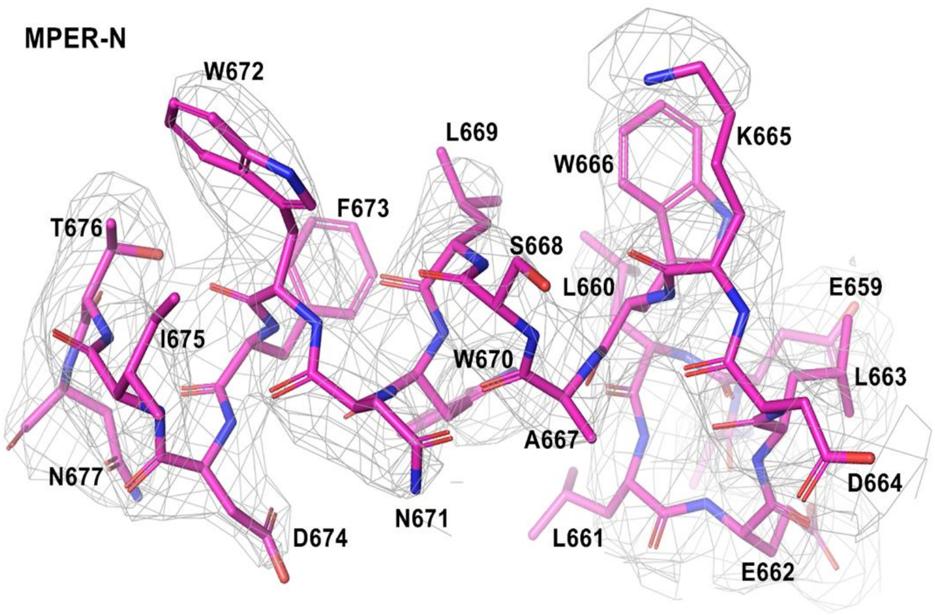


**Supplementary Figure 7. Electron density and model of the MPER-C in the crystal structure of the Fab460/MPER-C complex at a resolution limit of 3.6 Å.** The MPER-C peptide is drawn in stick representation. The grey mesh represents the σA-weighted $2F_o$-$F_c$ electron density map with a contour level of 1.0 σ.

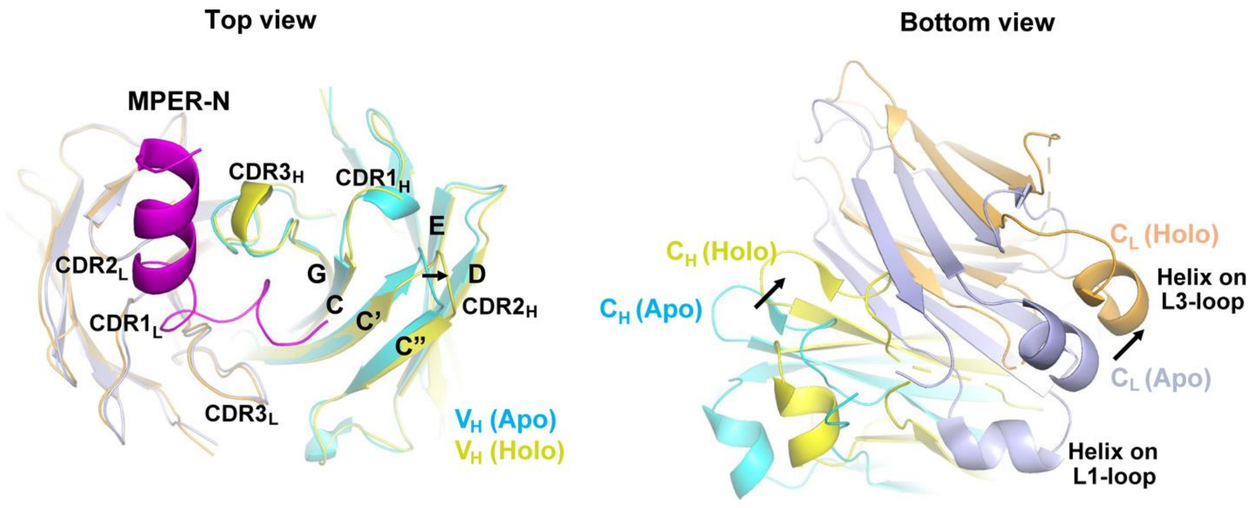


**Supplementary Figure 8**. **Conformational alterations in Fab460 upon MPER-N binding**. To measure the conformational change in the Fab after MPER-N binding, the $V_L$ domains from the structures of Fab460 alone (Apo) and in complex with MPER-N (Holo) were superimposed. The RMSD value obtained from the superposition is 0.45 Å (for all atoms). The left panel shows a top view of the aligned structures and illustrates that MPER-N binding results in a slight expansion of the Fab460 binding groove on the side of the C"-strand of the $V_H$ domain, represented by a relative shift of the $CDR2_H$ loop. The right panel shows a bottom view of the aligned structures and reveals that MPER-N binding results in a shift of the base of the $C_LC_{H1}$ domains, consequent to a change in the elbow angle.

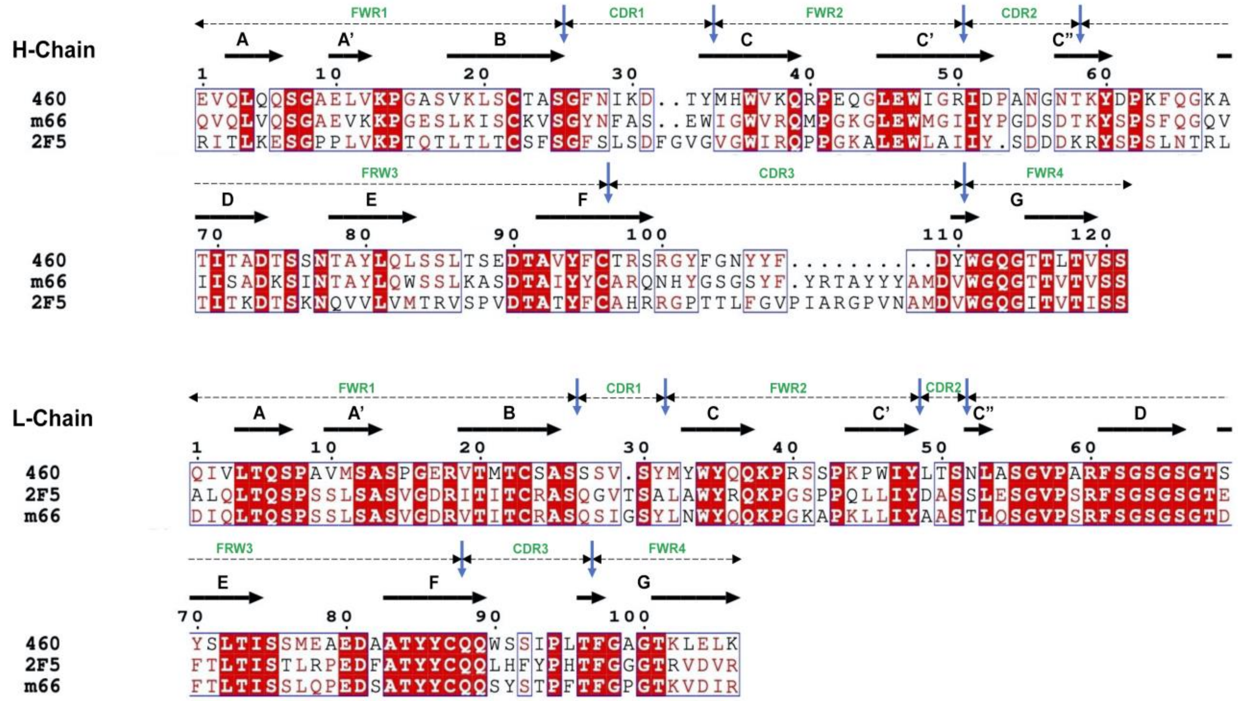


**Supplementary Figure 9. Structure-based sequence alignment of Fabs 460, 2F5 and m66.** β-strands are marked with arrows above the appropriate sequences of Fab460. There is a bulge (not shown) in the G-strand in both the H- and L-chain. The CDRs and FWRs are labeled in green. Identical residues are highlighted in red. The alignment was generated with the programs MultAlin (1) and ESPript (2) and was modified for clarity. Note that IGHV1-2*02, IGHD4-23*01 and IGHJ4*01 germline gene usages are for the V, D, and J segments of FAb460 $V_H$ domain, and the $V_L$ domain segments (V and J) for 460 are IGKV7-3*01 and IGKJ4*02

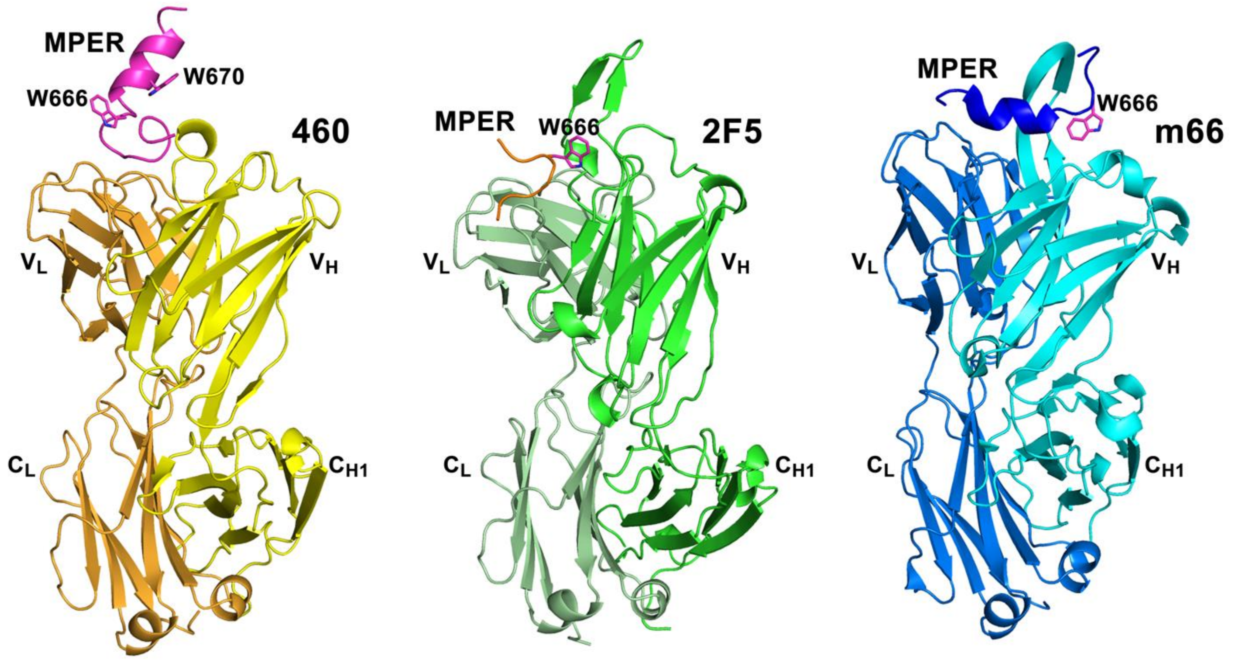


**Supplementary Figure 10. Ribbon diagrams of three Fabs targeting the MPER N-terminal region in complex with MPER peptides**. Fabs 460 (PDB: 8FZ2), 2F5 (PDB: 2FX7) and m66 (PDB: 4G6F) are displayed as ribbon diagrams and placed in a similar orientation. MPER residue W666 from their common β-turn is shown in stick representation to indicate the relative position of the structural motif. MPER residue W670 of the MPER N-terminal helix in the Fab460 complex is also shown in stick representation to indicate the helix orientation.

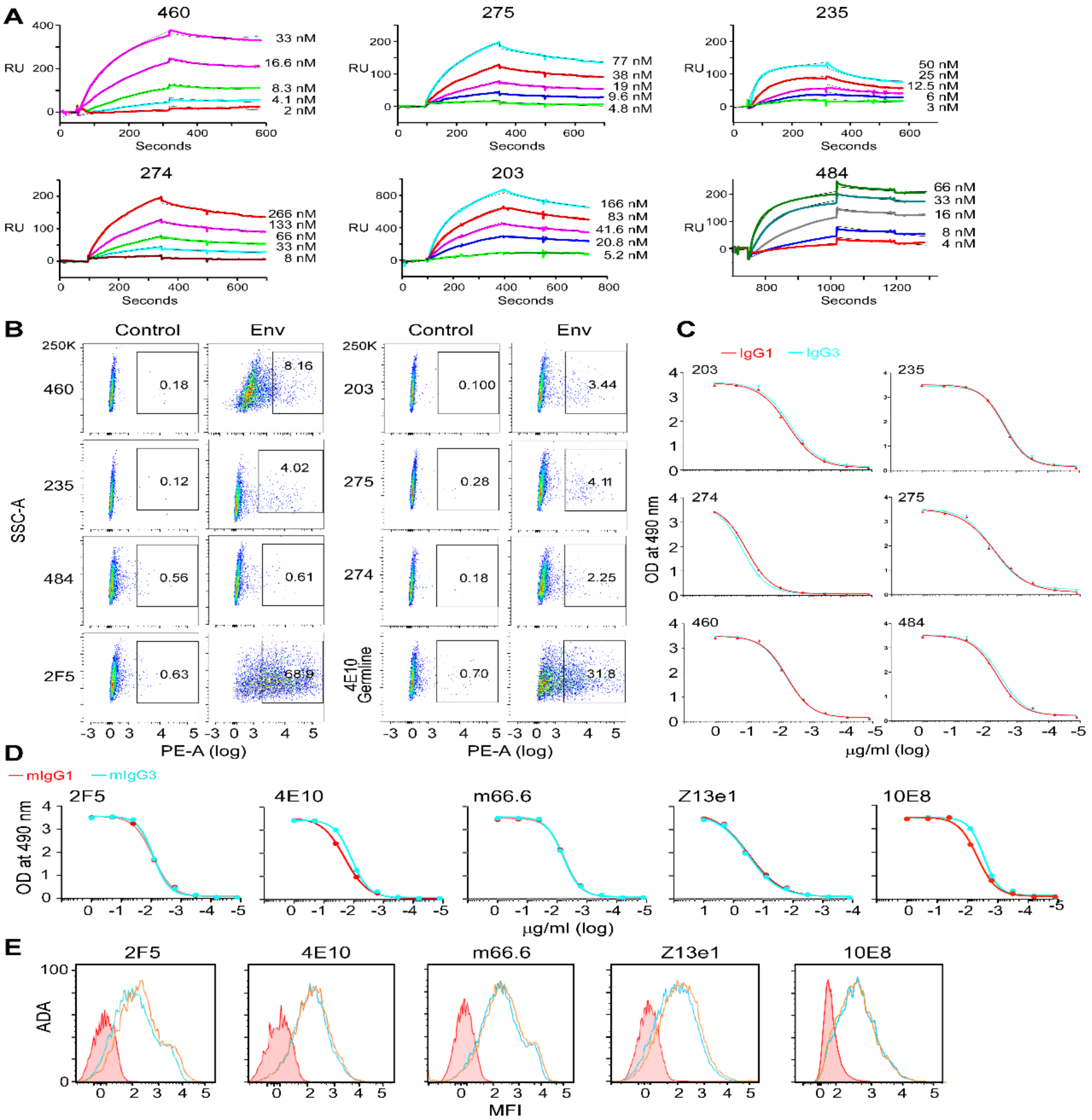


**Supplementary Figure 11. Characteristics in antigenicity between vaccine-elicited rmAbs vs human patient-derived bnAbs expressed as IgG1 and IgG3 subtypes.** (**A**) Biacore-binding kinetics of the indicated vaccine-elicited rmAbs to pMPER/liposome complexes (at a molar peptide:lipid ratio of 1:1000) captured on the surface of an L1 chip. Shown are binding sensograms of various rmAbs (color-coded at indicated concentrations) to pMPER/liposomes overlaid with fits of the data (black) in a 1:1 Langmuir model with baseline drift. (**B**) The flow cytometric analysis of the binding of the indicated Abs to 293T cells expressing ADA gp145 in comparison with un-transfected negative control cells and presented as dot plots. Cells were stained with Ab at 20 μg/ml for the vaccine-elicited 460, 235, 484, 203, 274 and 275 Abs or at 10 μg/ml for the mature 2F5 and germline 4E10 bnAbs, respectively. (**C**) Relative binding affinity of vaccine-elicited rmAbs expressed as IgG1 and IgG3 forms with the same specificity determined by ELISA against pMPER/liposome at a molar peptide:lipid ratio of 1:50. (**D**) Relative binding affinity of IgG1 and IgG3 versions of human bnAbs 2F5, 4E10 and 10E8, and nAbs m66.6 and Z13e1, respectively, determined by ELISA against pMPER/liposome (at a molar peptide:lipid ratio of 1:50). (**E**) The flow cytometric analysis of gp145 binding by bnAbs 2F5, 4E10 and 10E8, and nAbs m66.6 and Z13e1 at 10 μg/ml. Cyan indicates the IgG1 and orange indicates the IgG3 subtype of the bnAbs along with a negative control shown in red. Two independent experiments were performed.

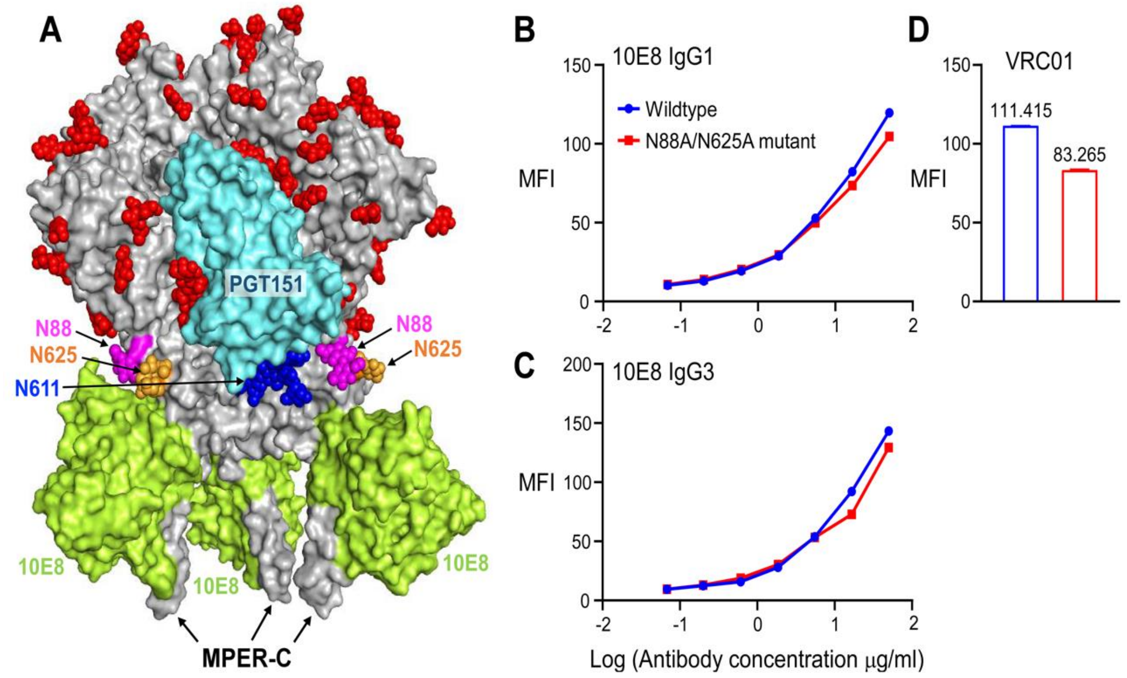


**Supplemememtary Figure 12. Effect of glycans proximal to the MPER of gp160 on anti-MPER bnAb 10E8 binding.** (**A**) A molecular surface representation of the trimeric HIV-1 envelope glycoprotein (in gray) in complex with one PGT151 Fab (in cyan) and three 10E8 Fabs (in limon). Glycans on the surface of the envelope glycoprotein are drawn in sphere representation. Three glycosylation sites (including glycans) close to MPER, N88, N611 and N625 are colored in magenta, blue and orange, respectively. More than one representation of each color is the result of the trimeric structure. 10E8 Fab binds to the MPER-C helix. This diagram is based on the structure of PDB 6VPX (3). (**B and C**) Concentration-dependent (log 10) binding by flow cytometry of 10E8 as IgG1(B) and IgG3 (C) subtypes to ADA gp145 (blue) and (**C**) ADA gp145 N88A/N625A mutant trimers (red). Two independent experiments were performed. (**D**) Expression levels of the wildtype and the glycan mutant gp145 as assessed by bnAb VRC01 binding at 10 μg/ml directed to the CD4 binding site.

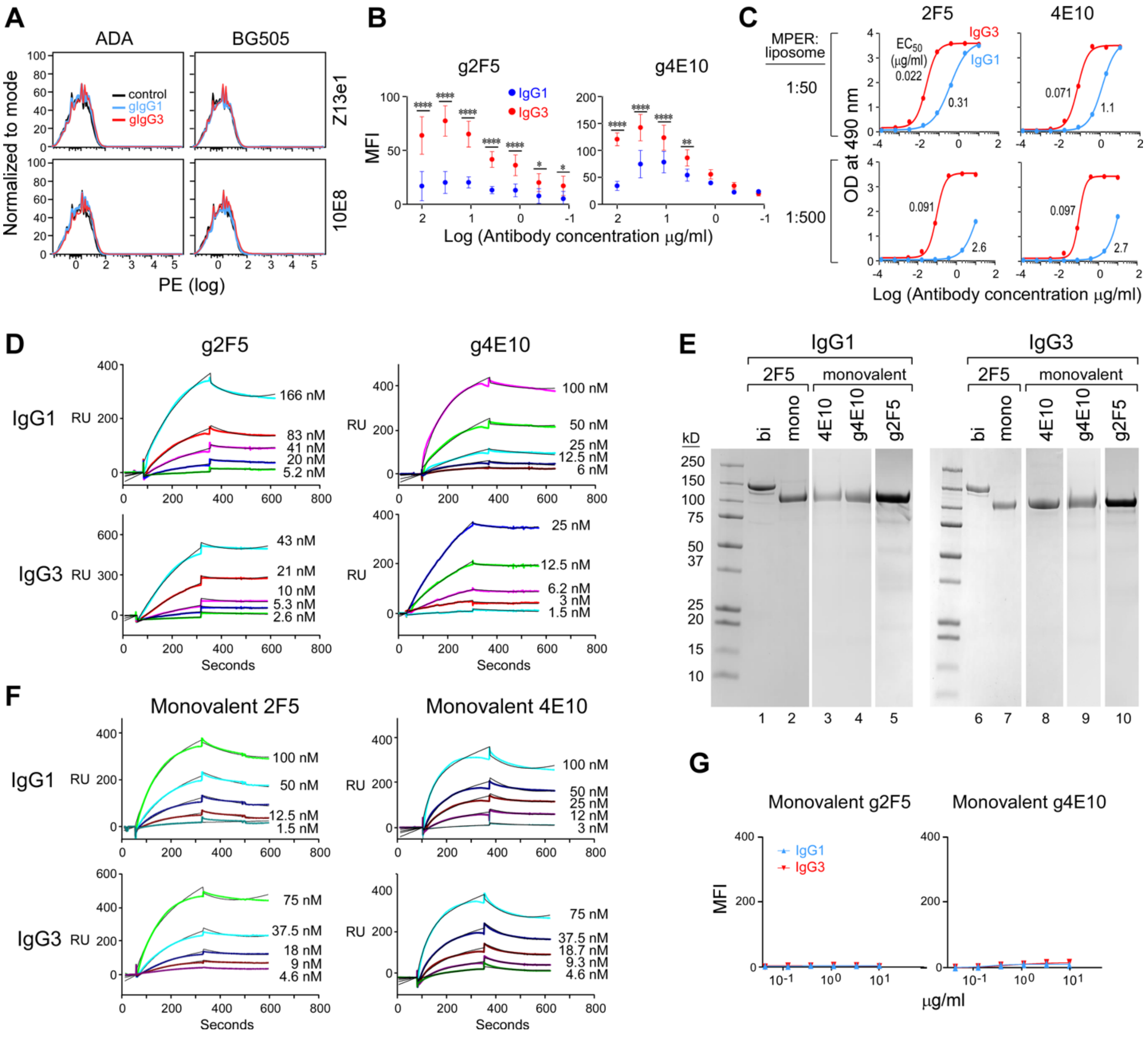


**Supplementary Figure 13. Composite binding analyses of intact germline 2F5 and 4E10, monovalent germline 2F5 and 4E10, and mature monovalent 2F5 and 4E10 bnAbs expressed both as IgG1 and IgG3 subtypes.** (**A**) Germline reverted 10E8 and Z13e1 binding to ADA and BG505 gp145 expressed on the surface of 293T cells. IgG1 and IgG3 forms of the antibodies were assessed for Env binding at 30 µg/ml indicated in cyan, orange and control cells in red, respectively. Two independent experiments were performed. (**B**) Concentration-dependent binding of germline-reverted, intact 2F5 and 4E10 as IgG1 and IgG3 forms for ADA gp145 in flow cytometry. Data are represented as mean of four independent experiments, respectively. Error bars represent standard deviations. Statistically significant differences between IgG subtypes were determined by 2-way ANOVA, denoted by p values: ****, p<0.0001; **,

$p<0.01$. (**C**) Relative binding affinity of germline 2F5 and germline 4E10 IgG1 (cyan) in comparison with that of the respective IgG3 subtype (red). The $EC_{50}$ of each antibody was determined by ELISA against pMPER/liposome at high (molar peptide:lipid ratio of 1:50) and low density of MPER peptide (molar peptide:lipid ratio of 1:500), respectively. (**D**) Biacore binding sensograms of germline 2F5 and germline 4E10 IgG1 and IgG3 subtypes to MPER/liposome complex captured on the surface of an L1 chip as detailed in Methods. (**E**) Nonreducing SDS-PAGE (8%) showing the migration of the indicated purified monovalent anti-MPER IgG1 antibodies as well as their respective monovalent IgG3 of the same specificity. Bivalent 2F5 IgG1 and IgG3 versions are shown for comparison. (**F**) Biacore binding kinetics of mature monovalent 2F5 and 4E10 IgG1 and IgG3 subtypes binding to pMPER/liposomes as detailed in Methods. (**G**) Concentration-dependent binding of germline-reverted monovalent 2F5 and 4E10 IgG1 in comparison with that of IgG3 with the same specificity for ADA gp145 when expressed on the 293T cell surface in flow cytometry. Representative of two independent experiments.

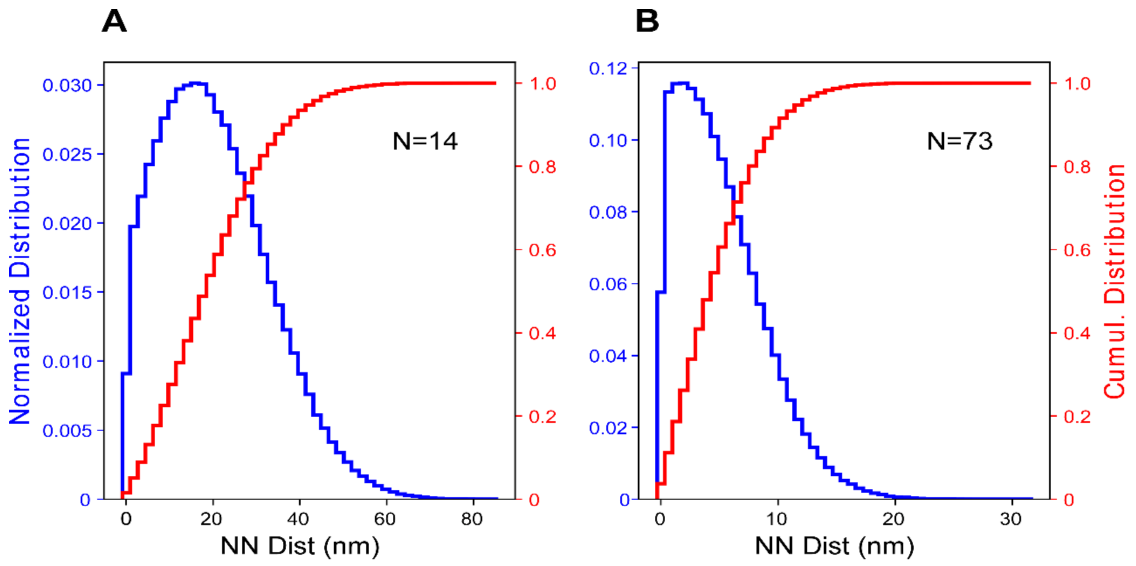


**Supplementary Figure 14**. **Normalized (blue) and cumulative (red) distribution of nearest-neighbor distances of spikes on virions.** (**A**) 14 spikes per virion, (**B**) 73 spikes per virion. Distances are measured between surfaces, not between centers of spikes, which is more relevant to crosslinking by two Fabs of an IgG. In each case, 50 bins were used to generate histograms. From the cumulative distributions for N=14, the probability that the nearest-neighbor distance is less than 10 nm is 0.23 and it is 0.48 for 17 nm. For N=73, the probability is already 0.90 for 10 nm. Thus, there is about 2-fold difference in the probability that a given virion has at least two spikes that can be crosslinked by IgG1 versus IgG3 on HIV-1 virions. This difference may be an underestimate, given that the shorter length and more limited flexibility of the IgG1 hinge relative to that of IgG3 may be less able to mediate inter-spike crosslinking when the distance between two spikes is small (4) and the approaching angle of two Fab arms are restricted by the geometry of the MPER between those spikes.

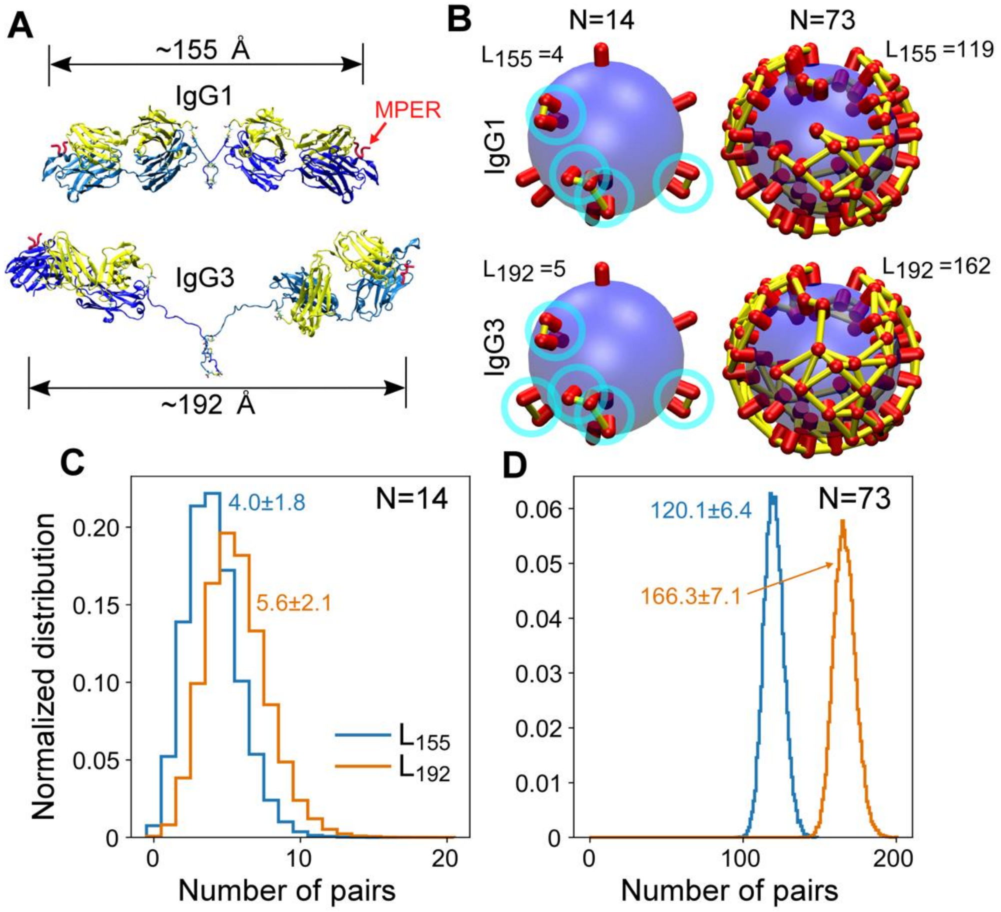


**Supplementary Figure 15. Comparison between IgG1 and IgG3 with greater spans**. See Fig. 6 for explanation of each panel. In **(A)**, orientations of Fab domains are arbitrarily set to provide an approximate estimate of their spans. In the case of the IgG1 model, the two Fab domains form a nearly 180º angle.

**Supplementary Table 1. Data collection and refinement statistics**

| ***Data collection*** | **235 Apo** | **235 + MPER-C** | **460** | **460 + MPER-N** |
|---|---|---|---|---|
| Space group | $P4_22_12$ | $P1$ | $P3_121$ | $H32$ |
| Unit Cell dimensions | | | | |
| *a, b,* c (Å) | 120.8, 120.8, 86.94 | 43.48, 108.3, 121.2 | 129.6, 129.6, 96.81 | 144.1, 144.1, 149.4 |
| *α, β, γ* (°) | 90, 90, 90 | 85.63, 82.95, 85.59 | 90, 90, 120 | 90, 90, 120 |
| Mol or Complex (AU) | 1 | 4 [2] | 1 | 1 |
| Wavelength (Å) | 0.9792 | 0.9792 | 0.9791 | 0.9792 |
| Resolution (Å) | 1.94 – 49.60 | 2.45 – 47.95 | 2.00 – 44.40 | 3.50 – 41.59 |
| Number of Unique | 46,097 (2,312) [1] | 78,049 (3,908) [3] | 63,553 (3,133) [4] | 7,498 (482) [5] |
| Completeness (%) | 96.7 (98.9) [1] | 96.3 (96.9) [3] | 99.9 (99.9) [4] | 97.5 (94.1) [5] |
| Redundancy | 4.4 (4.1) [1] | 2.9 (2.7) [3] | 6.8 (6.4) [4] | 6.7 (5.7) [5] |
| $R_{merge}$ | 0.126 (0.973) [1] | 0.143 (0.811) [3] | 0.102 (1.193) [4] | 0.231 (1.158) [5] |
| $R_{pim}$ | 0.065 (0.532) [1] | 0.097 (0.579) [3] | 0.041 (0.489) [4] | 0.094 (0.510) [5] |
| $CC_{1/2}$ | 0.985 (0.505) [1] | 0.974 (0.386) [3] | 0.992 (0.571) [4] | 0.966 (0.549) [5] |
| I/σ(I) | 15.5 (1.5) [1] | 7.0 (1.0) [3] | 23.2 (1.4) [4] | 8.7 (1.3) [5] |
| Wilson B-factors (Å$^2$) | 25.7 | 32.4 | 37.0 | 68.2 |
| ***Phasing*** [6] | | | | |
| ***Refinement*** | | | | |
| Resolution (Å) | 1.94 – 49.60 | 2.45 – 47.95 | 2.00 – 44.40 | 3.50 – 41.59 |
| $R_{work}/R_{free}$ | 17.1/20.6 | 20.0/24.4 | 16.7/19.4 | 26.2/31.0 |
| No.of atoms | | | | |
| Protein | 3,469 | 12,663 | 3,331 | 3,010 |
| Water/Others | 242/120 | 433/49 | 249/96 | 0/0 |
| B-factors (Å$^2$) | | | | |
| Protein atoms | 31.7 | 41.0 | 43.3 | 99.2 |
| Water/Others | 35.3/49.5 | 35.0/43.8 | 48.0/65.9 | NA/NA |
| R.m.s deviation | | | | |
| Bond length (Å) | 0.001 | 0.002 | 0.014 | 0.001 |
| Bond angle (°) | 1.042 | 0.484 | 1.286 | 0.470 |
| Ramachandran Plot (%) | | | | |
| Preferred regions | 97.74 | 97.15 | 97.69 | 92.36 |
| Allowed regions | 2.26 | 2.85 | 2.31 | 7.40 |
| Outliers | 0.00 | 0.00 | 0.00 | 0.24 |
| All-atom clashscore | 4.11 | 3.11 | 3.29 | 6.32 |
| Rotamer outliers (%) | 0.25 | 2.11 | 0.79 | 0.00 |
| $C_β$ deviations | 0.00 | 0.00 | 0.00 | 0.00 |
| **PDB code** | 8FWF | 8FYM | 8FXJ | 8FZ2 |

[1] (Last resolution bin, 1.94 – 1.97 Å); [2] Four complexes in one asymmetrical unit are distinguished from each other. There is no extra symmetry for a space group of higher symmetry; [3] (Last resolution bin, 2.45 – 2.49 Å); [4] (Last resolution bin, 2.00 – 2.03Å); [5] (Last resolution bin, 3.50 – 3.57 Å); [6] By molecular replacement method. Initial search templates for 235 and 460 *apo* structures are the $V_LV_H$-module and

$C_LC_{H1}$-module from structures of PDB codes 3GKW and 3BKY, respectively. Structural determinations were straightforward. Subseqeuntly, the *apo* structures were used for the structural determination of complexes of 235 and 460.

**Supplementary Table 2. Properties of the interface between 235 and MPER-C**

**Buried Surface Areas (Å$^2$)**

| Fab235/MPER-C complex | LHP | ABC | DEF | GIJ | Average |
|---|---|---|---|---|---|
| **Between H-chain and MPER-C** | 930.1 | 888.0 | 909.7 | 862.6 | **897.6** |
| **Between L-chain and MPER-C** | 101.6 | 99.7 | 158.2 | 155.8 | **128.8** |
| **Total** | 1031.7 | 987.7 | 1067.9 | 1018.4 | **1026.4** |

**Major specific interactions between Fab235 and MPER-C (bond length, Å)**
(Not including water-bridged interactions)

| Fab235/MPER-C complex | LHP | ABC | DEF | GIJ |
|---|---|---|---|---|
| **Contribution from Fab235 H-chain** | | | | |
| $Y107_H$ [OH] and MPER N674 [ND2] | 2.84 | 3.37 | 3.22 | 3.39 |
| $Y107_H$ [OH] and MPER N674 [OD1] | | 3.59 | | 3.26 |
| $N52_H$ [ND2] and MPER N677 [OD1] | 2.92 | 2.75 | 2.70 | 2.77 |
| $Y101_H$ [OH] and MPER N677 [OD1] | 2.51 | 2.44 | 2.62 | |
| $Y101_H$ [OH] and MPER-C Y681 [N] | 3.68 | 3.71 | 3.74 | 3.72 |
| **Contribution from Fab235 L-chain** | | | | |
| $N99_L$ [OD2] and MPER S688 [OG] | | | 3.40 | 2.91 |

The contents of these tables were generated with the online server program **PDBePISA** (Proteins, Interfaces, Structures and Assemblies). https://www.ebi.ac.uk/pdbe/pisa/

**Supplementary Table 3. Properties of the interface between Fab460 and MPER-N**

**Buried Surface Areas (Å$^2$)**

| **Fab460/MPER-N complex** | |
|---|---|
| **Between H-chain and MPER-N** | 757.9 |
| **Between L-chain and MPER-N** | 486.9 |
| **Total** | 1244.7 |

**Major specific interactions between Fab460 and MPER-N (bond length, Å)**
(No potential water-bridged interactions can be identified due to the low resolution limit)

| **Fab460/MPER-C Complex** | |
|---|---|
| **Contribution from Fab460 H-chain** | |
| $N57_H$ [OD1] and MPER K658 [N] | 3.55 |
| $K59_H$ [NZ] and MPER D659 [OD1] | 2.25 |
| $Y102_H$ [N] and MPER L661 [O] | 2.45 |
| $R50_H$ [NH1] and MPER E662 [OE1] | 2.31 |
| $R50_H$ [NH2] and MPER E662 [OE2] | 2.45 |
| $K59_H$ [NZ] and MPER E662 [OE2] | 2.51 |
| $Y106_H$ [OH] and MPER W666 [O] | 3.53 |
| $Y102_H$ [OH] and MPER N671 [OD1] | 2.70 |
| **Contribution from Fab460 L-chain** | |
| $W90_L$ [O] and MPER D664 [N] | 2.86 |
| $Y33_L$ [N] and MPER D664 [OD1] | 3.61 |
| $S91_L$ [N] and MPER D664 [OD2] | 3.16 |
| $S91_L$ [OG] and MPER D664 [OD2] | 3.08 |
| $Y31_L$ [OH] and MPER S668 [OG] | 3.61 |

The contents of these tables were generated with the online server program
**PDBePISA** (Proteins, Interfaces, Structures and Assemblies). https://www.ebi.ac.uk/pdbe/pisa/

**Supplementary Table 4. Neutralization potency of mature 2F5 and 4E10 IgG1 and IgG3 subtypes against 12 isolate HIV-1 Env pseudoviruses.**

| | $IC_{50}$ (mg/ml) | | | | $IC_{80}$ (mg/ml) | | | |
|---|---|---|---|---|---|---|---|---|
| | 2F5 IgG1 | 2F5 IgG3 | 4E10 IgG1 | 4E10 IgG3 | 2F5 IgG1 | 2F5 IgG3 | 4E10 IgG1 | 4E10 IgG3 |
| BB1056-10.TA11.1826 | 0.234 | 0.473 | 0.321 | 1.017 | 1.504 | 2.610 | 3.874 | 8.683 |
| 191955_A11 | 1.610 | 2.328 | 1.160 | 3.170 | 8.777 | 14.284 | 5.608 | 17.434 |
| C3347.c11 | 0.153 | 0.169 | 0.064 | 0.196 | 1.019 | 1.101 | 0.531 | 0.931 |
| QH0692.42 | 2.015 | 1.408 | 2.425 | 7.209 | 12.728 | 9.775 | 18.092 | 50.000 |
| SC422661.8 | 0.786 | 0.599 | 1.226 | 2.744 | 4.364 | 5.432 | 6.684 | 17.193 |
| REJO4541.67 | 0.403 | 0.630 | 0.356 | 1.707 | 4.421 | 6.484 | 3.801 | 14.324 |
| 6952.v1.c20 | 0.241 | 0.298 | 0.506 | 1.032 | 1.989 | 2.502 | 4.793 | 8.303 |
| WITO4160.33 | 0.512 | 0.537 | 0.232 | 0.556 | 3.549 | 5.455 | 1.574 | 4.891 |
| 231965.c1 | 20.532 | 42.852 | 34.667 | 50.000 | 50.000 | 50.000 | 50.000 | 50.000 |
| BG505/T332N | 0.299 | 0.142 | 0.362 | 0.971 | 1.325 | 0.824 | 1.757 | 4.817 |
| MuLV (Neg. Cont) | 50.000 | 50.000 | 50.000 | 50.000 | 50.000 | 50.000 | 50.000 | 50.000 |

Color scale: >50; 30-50; 20-30; 10-20; 1.0-10; 0.1-1.0; <0.1

**Supplementary Table 5. Comparison of the neutralization potency of the IgG1 and IgG3 subtypes of monovalent mature 2F5 and 4E10 against 12 isolate HIV-1 Env pseudoviruses.**

| | $IC_{50}$ (mg/ml) | | | | $IC_{80}$ (mg/ml) | | | |
|---|---|---|---|---|---|---|---|---|
| | monovalent 4E10 IgG1 | monovalent 4E10 IgG3 | monovalent 2F5 IgG1 | monovalent 2F5 IgG3 | monovalent 4E10 IgG1 | monovalent 4E10 IgG3 | monovalent 2F5 IgG1 | monovalent 2F5 IgG3 |
| BB1056-10.TA11.1826 | 3.328 | 3.173 | 6.286 | 5.896 | 22.413 | 27.325 | 21.341 | 31.703 |
| 191955_A11 | 8.879 | 18.250 | 16.220 | 27.090 | 30.534 | 50.000 | 50.000 | 50.000 |
| C3347.c11 | 4.496 | 1.386 | 11.210 | 4.857 | 21.746 | 16.132 | 33.562 | 25.457 |
| QH0692.42 | 42.790 | 47.370 | 15.230 | 15.570 | 50.000 | 50.000 | 50.000 | 50.000 |
| SC422661.8 | 16.140 | 16.990 | 7.863 | 8.809 | 48.797 | 50.000 | 25.656 | 47.418 |
| REJO4541.67 | 4.446 | 2.937 | 6.114 | 5.158 | 33.594 | 40.534 | 24.861 | 33.281 |
| 6952.v1.c20 | 4.158 | 4.927 | 1.986 | 2.497 | 23.243 | 33.418 | 8.266 | 10.825 |
| WITO4160.33 | 3.020 | 2.585 | 10.500 | 9.155 | 24.949 | 22.904 | 35.102 | 50.000 |
| 231965.c1 | 50.000 | 50.000 | 50.000 | 50.000 | 50.000 | 50.000 | 50.000 | 50.000 |
| BG505/T332N | 50.000 | 6.001 | 21.690 | 5.555 | 21.244 | 23.993 | 50.000 | 42.032 |
| MuLV (Neg. Cont) | 50.000 | 50.000 | 50.000 | 50.000 | 50.000 | 50.000 | 50.000 | 50.000 |

>50 | 30-50 | 20-30 | 15-20 | 5-15 | <5
